# Structural network maturation of the preterm human brain

**DOI:** 10.1101/195800

**Authors:** Tengda Zhao, Virendra Mishra, Tina Jeon, Minhui Ouyang, Qinmu Peng, Lina Chalak, Jessica Lee Wisnowski, Roy Heyne, Nancy Rollins, Ni Shu, Hao Huang

**Affiliations:** State Key Laboratory of Cognitive Neuroscience and Learning & IDG/McGovern Institute for Brain Research, Beijing Normal University, Beijing, China 100875; Center for Collaboration and Innovation in Brain and Learning Sciences, Beijing Normal University, Beijing, China 100875; Beijing Key Laboratory of Brain Imaging and Connectomics, Beijing Normal University, Beijing100875, P. R. China; Advanced Imaging Research Center, University of Texas Southwestern Medical Center, Dallas, TX 75390; Radiology Research, Children’s Hospital of Philadelphia, Philadelphia, PA 19104; Department of Pediatrics, University of Texas Southwestern Medical Center, Dallas, TX 75390; Department of Radiology, University of Texas Southwestern Medical Center, Dallas, TX 75390; Department of Radiology, Perelman School of Medicine, University of Pennsylvania, PA 19104; Department of Radiology, Children’s Hospital Los Angeles, Keck School of Medicine, University of Southern California, Los Angeles, CA

**Keywords:** brain maturation, baby connectome, structural connectivity, segregation, diffusion MRI

## Abstract

During the 3^rd^ trimester, large-scale of neural circuits are formed in the human brain, resulting in the adult-like brain networks at birth. However, how the brain circuits develop into a highly efficient and segregated connectome during this period is unknown. We hypothesized that faster increases of connectivity efficiency and strength at the brain hubs and rich-club are critical for emergence of an efficient and segregated brain connectome. Here, using high resolution diffusion MRI of 77 preterm-born and term-born neonates scanned at 31-42 postmenstrual weeks (PMW), we constructed the structural connectivity matrices and performed graph-theory-based analyses. We found faster increases of nodal efficiency mainly at the brain hubs, distributed in primary sensorimotor regions, superior-middle frontal and posterior cingulate gyrus during 31-42PMW. The rich-club and within-module connections were characterized by higher rates of edge strength increases. Edge strength of short-range connections increased faster than that of long-range connections. The nodal efficiencies of the hubs predicted individual postmenstrual ages more accurately than those of non-hubs. Collectively, these findings revealed regionally differentiated maturation in the baby brain structural connectome and more rapid increases of the hub and rich-club connections, which underlie network segregation and differentiated brain function emergence.

## Introduction

In the last a few weeks prior to normal time of birth, a large scale of brain circuit formation underlies the emergence of human connectome at a macroscale. Massive development of cortico-cortical axonal pathways before birth offers structural basis to establish an adult-like brain network (e.g. Kostović and Jovanov-Milošević 2006; Kostović and Judaš 2010). The neuronal activities associated with the circuit formation undergo substantial remodeling after birth (e.g. LaMantia and Rakic 1990; Innocenti and Price 2005). As suggested by previous neuropathological studies (e.g. Huttenlocher and Dabholkar 1997), regionally differentiated developments of neuronal connections are associated with heterogeneous emergence of brain functions, with primary sensorimotor function generally emerging earlier than higher-order cognitive functions. However, little is known about the structural organization of neural networks at the macroscale during this critical period. Knowledge of the ontogeny of the human connectome during late fetal development may provide not only insight into normal brain development, but also a reference for elucidating the complex trajectories of atypical or abberant neurodevelopment.

The structural connections have recently been extensively studied by magnetic resonance imaging (MRI), capable of surveying entire brain connectivity noninvasively. Diffusion MRI (dMRI), a type of MRI methods, has been applied as an approach to infer axonal pathways constituting the structural brain connectivity *in vivo*. With dMRI-based tractography (e.g. Mori et al. 1999), the emergence of brain white matter (WM) fibers has been delineated in the fetal brain as early as the beginning of 2^nd^ trimester (e.g. Huang et al. 2006; Huang et al. 2009; Vasung et al. 2010; Takahashi et al. 2012; Ouyang et al. 2015), consistent to the histological atlases (Bayer and Altman 2004). These dMRI studies have demonstrated, for example, that limbic WM fibers appear earlier while the association WM fibers constituting major cortico-cortical connectivity appear later. By the start of the 3^rd^ trimester, except arcuate fasciculus, all major WM fibers can be identified with dMRI (Feng et al., 2016). The asynchronous and heterogeneous maturation of WM across regions in the 3^rd^ trimester has also been suggested by other neuroimaging studies (e.g. Hüppi et al. 1998; Partridge et al. 2004; Bui et al. 2006; Aeby et al. 2009). In addition, it has been found that different cortical regions undergo differential maturation pattern in terms of cortical microstructure (McKinstry et al. 2002; DeIpolyi et al. 2005; Huang et al. 2013; Yu et al. 2016), also assessed with dMRI.

Although the literature above revealed spatiotemporally heterogeneous development of both cortical regions and WM pathways linking them, few studies have delineated the differential maturation pattern of structural connectivity from the perspective of a macro-scale connectome during the 3^rd^ trimester. The baby brain connectome (For a review, see e.g. Cao et al. 2017b) reveals the inter-regional connectivity pattern, in contrast to individual WM fiber bundles or brain regions. In a brain connectome, some regions are more interconnected with other brain regions, constituting “hubs” within the global network topography (e.g. Hagmann et al. 2008; Gong et al. 2009). Further, these hub regions tend to be densely interconnected with each other forming a rich-club organization (van den Heuvel and Sporns 2011), which serves as a highly efficient backbone for integration of neuronal activity across distributed circuits and presumably forms the foundation of complex neurological functions (van den Heuvel et al. 2012). With the structural connectivity underlying functional connectivity, the present connectomic study offers a unique view of understanding the structural substrate of emerging brain functions. The preterm and term-born brain connectome has been investigated with dMRI tractography and subsequent graph-theory analysis (Tymofiyeva et al. 2013; Ball et al. 2014; Brown et al. 2014; van den Heuvel et al. 2015; Batalle et al. 2017). These studies support the emergence of the hub regions and rich club organization during the 3^rd^ trimester (Ball et al. 2014; van den Heuvel et al. 2015). However, regionally differential maturational rates during the 3^rd^ trimester quantified by connectomic measures of brain hubs, rich-club and modules as well as short-range and long-range connections have not been determined. In addition, it remains to be determined how differentiated connectional maturation contributes to the segregation process of structural organization of baby brain.

In this study, we hypothesized that differentiated maturation of structural connectivity across brain regions plays a central role in emergence of an efficient and segregated brain connectome at birth. Relatively high resolution (1.5 × 1.5 × 1.6 mm^3^) dMRI images of 77 preterm-born or term-born neonates scanned around 31 to 42 postmenstrual weeks (PMW) (Engle 2004) were acquired. The structural connectivity matrix of each neonate was constructed with dMRI tractography. With the comprehensive graph theory analysis at global, modular and regional connection levels, we examined cross-sectional age-dependent developmental rate of the preterm and term-born brain network measures across different brain regions and connections.

## Materials and Methods

### Preterm subjects

The study was approved by the Institutional Review Board (IRB) of the University of Texas Southwestern Medical Center. 77 normal neonates (47 males and 30 females) were recruited from Parkland Memorial Hospital at Dallas. These neonates were scanned between 31.9 to 41.7 PMW, with postmenstrual age defined in accordance with Engle’s criteria (Engle 2004). All neonates underwent MR imaging as part of a study of normal prenatal and perinatal development; no neonates were scanned under clinical indications. Moreover, these neonates were recruited after rigorous screening procedures conducted by a board-certified neonatologist (LC) and an experienced pediatric radiologist (NR), based on subjects’ ultrasound, clinical MRI and medical record of the neonates and their mothers. Exclusion criteria include evidence of bleeding or intracranial abnormality by serial sonography; mother’s excessive drug or alcohol abuse during pregnancy; grade III-IV intraventricular hemorrhage; periventricular leukomalacia; hypoxic-ischemic encephalopathy; lung disease or brochopulmonary dysplasia; body or heart malformations; chromosomal abnormalities; necrotizing enterocolitis that requires intestinal resection or complex feeding/nutritional disorders; defects or anomalies of forebrain, brainstem or cerebellum; brain tissue dys-or hypoplasias; abnormal meninges; alterations in the pial or ventricular surface; or white matter lesions. Written and informed parental consents were obtained from the subject’s mother (or father if married). Detailed characteristics regarding this cohort are provided in Table 1.

**Table 1.** Demographic information of scanned neonates (^a^C for C-section and V for vaginal birth; ^b^B for breast-feeding and F for formula).

|  | Number of infants | Age range (weeks) | Age mean (weeks) | Weight range (kg) | Weight mean (kg) | Male, n (%) | White, n (%) | Mode of delivery | Feeding practice | Antibiotic exposure during pregnancy |
| --- | --- | --- | --- | --- | --- | --- | --- | --- | --- | --- |
| **At birth** | 77 | 25.0-41.4 | 33.7 | 0.8-4.0 | 2.1 | 47 (61) | 59 (77) | C:29; V:48 | B: 77; F: 0 | Yes |
| **At scan** | 77 | 31.9-41.7 | 37.2 | 1.4-4.1 | 2.6 | 47 (61) | 59 (77) | C:29; V:48 | B: 77; F: 0 | Yes |

### MRI Acquisition

All neonates were scanned with a Philips 3.0 T Achieva MR scanner at the Children’s Medical Center, Dallas. They were well-fed before scanning. During scan, all neonates were asleep naturally without sedation. Earplugs, earphones and extra foam padding were applied to reduce the sound of the scanner while the neonates were asleep. A single-shot EPI sequence (SENSE factor = 2.5) was used for dMRI acquisition, with the following parameters: TE=78ms, TR=6850ms, in-plane field of view = 168 × 168mm^2^, in-plane imaging matrix = 112 × 112, in-plane imaging resolution =1.5 × 1.5mm^2^, slice thickness =1.6mm without gap, slice number=60, 30 independent diffusion encoding directions with b value = 1000 s/mm^2^. The images were reconstructed to 256 × 256 in-plane matrix. Two repetitions were conducted for dMRI acquisition, resulting in scan time of 11 minutes. As described in our previous publication (Huang et al. 2015), with 30 diffusion weighted image (DWI) volumes and 2 repetitions, we accepted those dMRI datasets with less than 5 DWI volumes affected by severe motion. The affected volumes were replaced by the good volumes of another dMRI repetition during postprocessing.

### Data preprocessing

Small motion and eddy current of dMRI of each neonate were corrected by registering all the DWIs to the b0 image using a 12-parameter (affine) automated image registration (AIR) algorithm (Woods et al. 1998). After AIR, six independent elements of the 3×3 diffusion tensor were determined by multivariate least-square fitting of DWIs (Basser et al. 1994). The tensor was diagonalized to obtain three eigenvalues (*λ*_1–3_) and eigenvectors (*V*_1–3_). Then the diffusion metrics, such as fractional anisotropy (FA) and apparent diffusion coefficient (ADC) images were calculated. All above-mentioned procedures were conducted offline using DTIStudio (Jiang et al. 2006).

### Network construction

Nodes and edges, the two fundamental elements of a network, were defined using the following procedures to construct the individual structural network.

#### Network node definition.

The nodes of each subject in the native dMRI space were obtained by transferring the parcellated cortical regions in the Johns Hopkins University (JHU) neonate atlas (Oishi et al. 2011). The contrasts of the single-subject b0 (ss-b0) image in the JHU atlas space (Fig. 1D) and individual neonate subject’s b0 image in the native space (Fig. 1A) were used to drive the nonlinear registration that transfers JHU atlas cortical parcellation to the individual neonate subjects. Briefly, neonate b0 image in the native space was registered to the ss-b0 images in the JHU atlas space with transformation T(•). The inverse transformation T^-1^(•) was used to map JHU atlas labels (Fig. 1E) to the native space of individual neonate (Fig. 1F). Discrete labeling values were preserved using a nearest-neighbor interpolation. The structural network (Fig. 1G) of each neonate was constructed with 58 cortical regions (Fig. 1F) representing 58 nodes of the brain network. The registration procedures were conducted using SPM8 software (http://www.fil.ion.ucl.ac.uk/spm/). Of note, the cortical regions were dilated by 9 voxels in order to allow traced WM fibers (see Network edge definition below) to reach the cortical nodes. To minimize spurious structural connections between the nodes, the voxels in the dilated cortical regions with ADC greater than 1.9 × 10^-3^mm^2^/s were likely to be those of cerebrospinal fluid (CSF) and removed.

**Figure 1.**
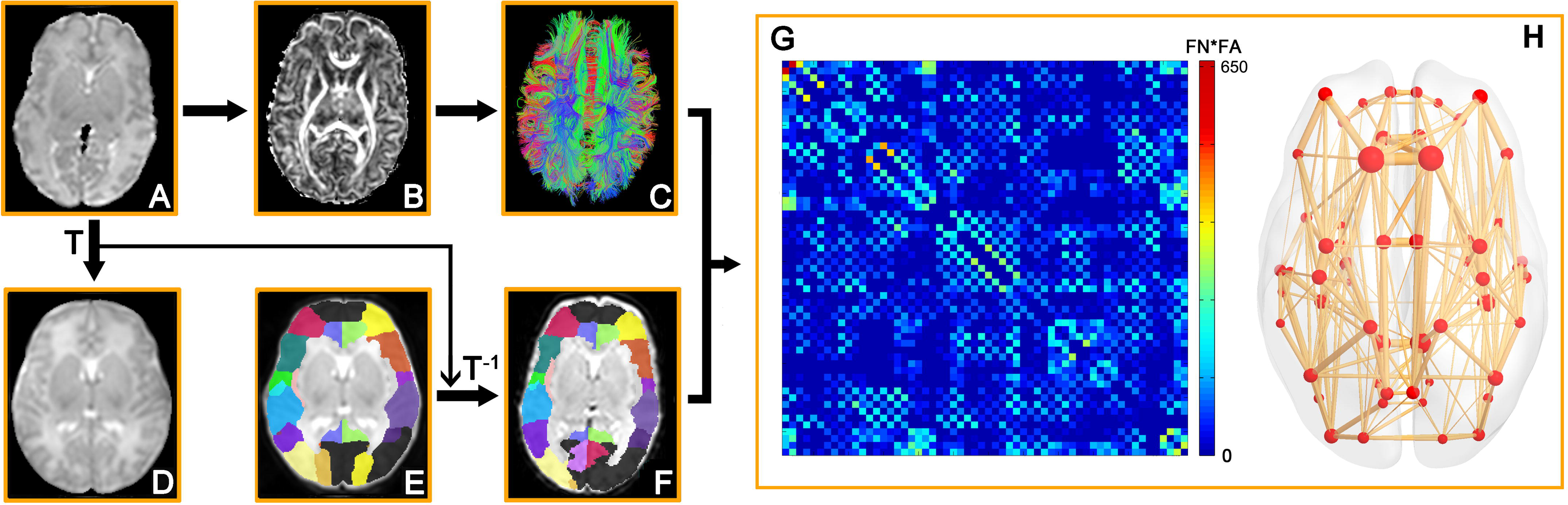
The flowchart of brain network construction. The b0 image of each neonate subject in its native space (A) was registered to the b0 image of the single-subject template in the JHU atlas space (D) with the transformation T(•). (B) and (C) are FA map and dMRI tractography results in the native space, respectively. The JHU atlas labels (E) were inversely transferred to the native space (F) with the transformation T^-1^(•). With delineation of network edges (C) and nodes (F) in the native space, the connectivity matrix (G) and network graph (H) were established. The flowchart was drawn demonstrating analysis of a representative neonate dataset. The reconstructed whole-brain fiber streamlines (C) and 3D representation of the structural network (H) were generated using TrackVis (http://trackvis.org/) and BrainNet Viewer software (Xia et al. 2013), respectively.

#### Network edge definition.

Network edges were defined with reconstructed WM fibers by dMRI tractography. Brute-force deterministic fiber tractography in the whole brain was performed with Diffusion Toolkit (http://trackvis.org/) (Mori et al. 1999; Huang et al. 2004). Due to low FA in preterm brains, the FA threshold was 0 (no FA threshold) (Takahashi et al., 2012) and angle threshold was 35^o^ for tractography. Only reconstructed streamlines with two end points located in the dilated cortical ribbon were kept, as shown in Fig. 1C. Two regions were considered structurally connected if there exists at least one streamline with two end-points located in these two regions. Anisotropy diffusivity such as FA and the streamlines obtained by tractography have both been proved as good markers for characterization of tissue microstructure and WM changes during development (Wimberger et al. 1995; Drobyshevsky et al. 2005; Huang et al. 2006; Huang et al. 2009; Takahashi et al. 2012). Therefore, we defined the number of fiber streamlines multiplied by the mean FA (FN × FA) of all connected fibers between two regions as the edge weight. As a result, we constructed a weighted structural network (Fig. 1H) for each neonate, represented by a symmetric 58 × 58 connectivity matrix (Fig. 1G).

### Network analysis

To describe the topological organization of the neonatal structural connectome, the following graph metrics were estimated, with detailed definitions of the network metrics provided in the Supplement material.

#### Global network organization.

For the global network metrics, we quantified the network sparsity, network strength (S_p_), global efficiency (E_glob_), local efficiency (E_loc_), shortest path length (L_p_), clustering coefficient (C_p_) small-world parameters (λ, γ and σ) (Rubinov and Sporns 2010).

#### Regional network efficiency.

To determine the nodal (regional) characteristics of the brain networks, we computed the nodal efficiency which is defined as (Achard and Bullmore 2007).

#### Hub distribution.

To identify the hub regions of the neonate connectome, we constructed the group-based backbone network by detecting the significant nonzero connections across all participants, with a nonparametric one-tailed sign test (p < 0.05, corrected) and assigning the edge weight with the group-averaged one. Based on the group-averaged backbone network, we identified the hub regions by sorting the nodal efficiency (E_nodal_(i) > mean+0.5*std). Next, the rich-club coefficient (*ϕ*) and normalized rich-club (RC) coefficient (*ϕ* _norm_) were calculated for the backbone network based on averaged network of all neonates, according to van den Heuvel and Sporns (2011). On the basis of the categorization of the nodes of the network into hub and non-hub regions, edges of the network were classified into. Finally, the rich-club, feeder and local edge strength was averaged edge weight of rich-club, feeder and local connections, respectively.

#### Short- and long-range edges.

For the reconstructed fibers, length was defined as the physical length of the streamline obtained by tractography (Fig. 6A) and the average physical length of all streamlines connecting each pair of brain regions was defined as the length of each connection. Then, the connections of each individual network were grouped into long-range and short-range ones based on the length of these connections. Considering the increasing brain size with age, we did not use a constant length threshold. Individual connections with length smaller/greater than average length of all connections were defined as short-/long-range connections, respectively.

#### Modular parcellation.

Module detection was performed with an optimized simulated annealing approach (Guimera et al. 2004) to parcellate the brain network into different modules (Newman and Girvan 2004). Briefly, the aim of this module identification process is to find a specific partition (p) which yields the largest network modularity, Q(p). Q(p) quantifies the difference between the number of intra-module links of actual network and that of random network in which connections are linked at random. The modular parcellation was performed on the individual network of each neonate brain with the modularity and module number of the individual network calculated. In addition, to apply a consistent modular parcellation across subjects, we also performed module detection on the backbone network. Based on the modular parcellation of backbone network, the participation coefficient (PC) of each node was calculated (Guimera et al. 2004; Sporns et al. 2007; Rubinov and Sporns 2010) to assess the contribution of each node to modular segregation or integration. Then, the hub regions were categorized as connector hubs (with PC > 0.5) which occupied high inter-module connections and provincial hubs (with PC < 0.5) which occupied high intra-module connections. Based on the backbone modular parcellation, the whole within- and between-module edge strength was the summation of edge weights of within- and between-module connections, respectively.

All network analyses were performed using GRETNA software (http://www.nitrc.org/projects/gretna/) (Wang et al. 2015) and the results were visualized using BrainNet Viewer software (https://www.nitrc.org/projects/bnv/) (Xia et al. 2013).

### Statistical analysis

#### Age effects on network properties.

To examine the age effects on the network topological properties, a general linear model (GLM) analysis was implemented between each network metric and postmenstrual age across all subjects, with gender and total brain volume (TBV) as covariates:

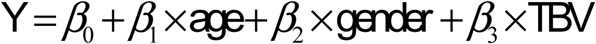

The slope of each metric against age *β*_1_ was used to represent the developmental rate.

#### λNetwork-based statistic (NBS).

To identify structural connections showing significant age effects from the whole connectome, we used the NBS approach (Zalesky et al. 2010). First, the same GLM analysis with gender and TBV as covariates was applied for the entire connectome. A threshold of p < 0.01 was used to yield t statistic matrix of suprathreshold connections. After that, the nonparametric NBS approach was used for controlling family wise error (FWE). By generate 5000 times permutation null distribution, the statistical significance of the observed component sizes in the un-corrected connection matrix was evaluated. Finally, the interconnected sub-network components with a corrected p < 0.01 were considered statistically significant.

### Clustering analysis

To group the brain regions with similar developmental trajectories, we used a data-driven k-means clustering method (Seber 2009). The set of brain regions with significant age-dependent nodal efficiency increase was used as the input and the developmental trajectory of each region’s efficiency was used as the feature of clustering. The k-means algorithm was initialized with randomized estimates for the trajectory centers and iterated multiple times to convergence. In our study, ten repetitions with different random initial cluster centroids were used to minimize the effect of start condition. The whole process was repeated with varying numbers of clusters from two to six and the final number of clusters was determined by the clustering results with the largest average silhouette value (Rousseeuw 1987). To test if nodal efficiency increases faster in the hubs than non-hubs during 32-41PMW, a two-cluster model with the highest silhouette value among the priori designs (2~6 clusters) was adopted.

### Prediction of the neonate age using support vector regression

A support vector regression (SVR) with a linear kernel function was used to test the prediction power of nodal efficiency on individual neonate postemenstrual age. The default settings with C = 1 and epsilon = 0.001 in the LIBSVM Toolbox (http://www.csie.ntu.edu.tw/~cjlin/libsvm/) were used to evaluate the SVR model (Dosenbach et al. 2010; Iuculano et al. 2014). A Leave-one-out cross-validation (LOOCV) was used to evaluate the prediction accuracy of the model. Each neonate was designated as the test data in turns while the remaining ones were used to train the SVR predictor which aimed at making a prediction about the test neonate’s age. Pearson correlation coefficient between the actual and predicted ages was calculated to assess the prediction accuracy. The nodal efficiencies of all regions or only hub/non-hub regions defined based on the networks of train samples were used as features for the SVR predictor separately. In each iteration, the hub distribution obtained from train sample was similar to that obtained from the whole cohort.

### Evaluation of the effects of different parcellation schemes

To evaluate the potential effects of the parcellation schemes (e.g. Zalesky et al., 2010) on the results, the neonate cortex was further randomly subdivided into 256 nodes with equal size to examine the age-dependent network property changes with a high-resolution parcellation. For each neonate, a high-resolution structural connectivity matrix in 256 by 256 was constructed. Same network analysis procedures and statistical analyses as those used in the low-resolution networks (58 nodes) were repeated.

## Results

### Age-dependent changes of global topological properties of neonate structural connectome

The global structural networks became stronger and more efficient from 32 to 41 PMW. About 5-fold increase of network strength (r = 0.46, p = 3.3 × 10^-5^), 6-fold increase of global efficiency (r= 0.46, p = 2.9 × 10^-5^) and 7-fold increase of local efficiency (r = 0.48, p = 1.1 × 10^-5^) were found (Fig. 2, upper panel). Prominent small-world organization was observed with *λ* ≈1 and *γ*>1 for the structural networks of all neonates aged 32 to 41 PMW. However, no significant age-dependent changes were found in number of edges (network sparsity) and small-worldness *σ* (p > 0.05, Fig. 2, lower panel). Age-dependent changes of other network measurements can be found in Supplemental Figure S1.

**Figure 2.**
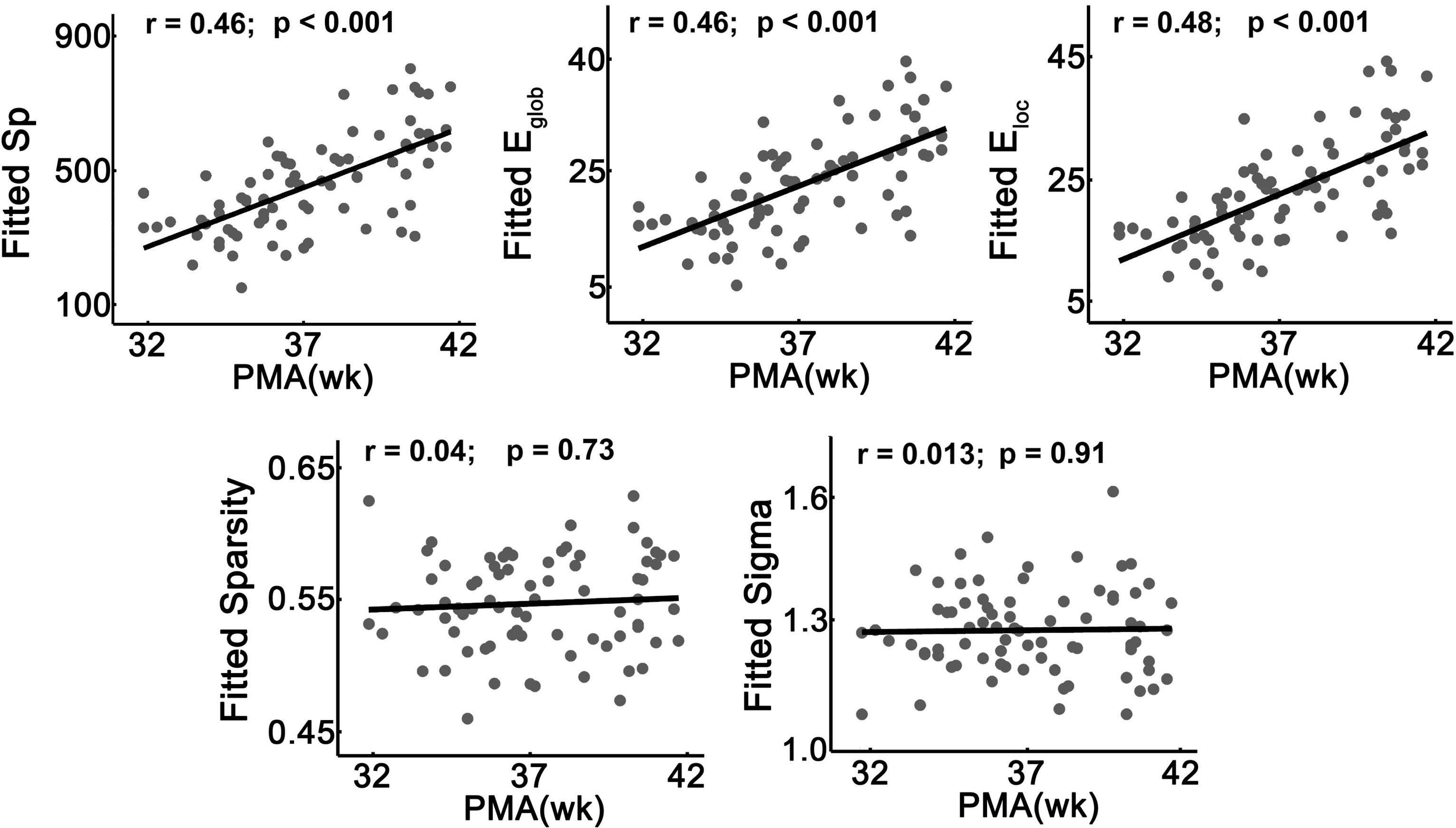
Age-related changes in the global network metrics of the neonate connectome. Significant age-related linear increases of the network strength, global efficiency and local efficiency (top panels) and non-significant age-related changes of sparsity and small-worldness (bottom panels) are demonstrated in the scatter plots.

### Differential nodal efficiency increases with faster nodal efficiency increases at the brain hubs

#### Differential nodal efficiency increases across brain regions

The heterogeneous distribution of nodal efficiency across brain regions is clear as shown by the fitted nodal efficiency maps of each week during 32-41PMW (Fig. 3A, left panel), with higher efficiency in prefrontal cortex, precentral and postcentral gyrus and lower efficiency in occipital cortex. Both the mean and the standard deviation (SD) of the nodal efficiency increased significantly with age (Mean: r = 0.46, p = 2.9 × 10^-5^; SD: r = 0.39, p = 6.1× 10^-4^) (Fig. 3A, right panel), indicating that both the average and the variability of nodal efficiency across the brain regions increased with age. As shown in Fig. 3B, 31 cortical regions widely distributed all over the brain in bilateral frontal, parietal, temporal and limbic regions exhibited significant age-related linear increases (p < 0.05, Bonferroni corrected) during 31-42PMW with GLM analysis. Importantly, the developmental rates of nodal efficiency varied across regions. More rapid, age-dependent increases were found in the precentral and postcentral gyrus, superior and middle frontal gyrus, precuneus and posterior cingulate gyrus relative to other brain regions (Fig. 3B). The scatter plots of three representative regions (left precentral gyrus: PrCG.L, left angular gyrus: ANG.L, and right parahippocampal gyrus: PHG.R) with distinguished nodal efficiency increase rates are shown in the right panel of Figure 3B.

**Figure 3.**
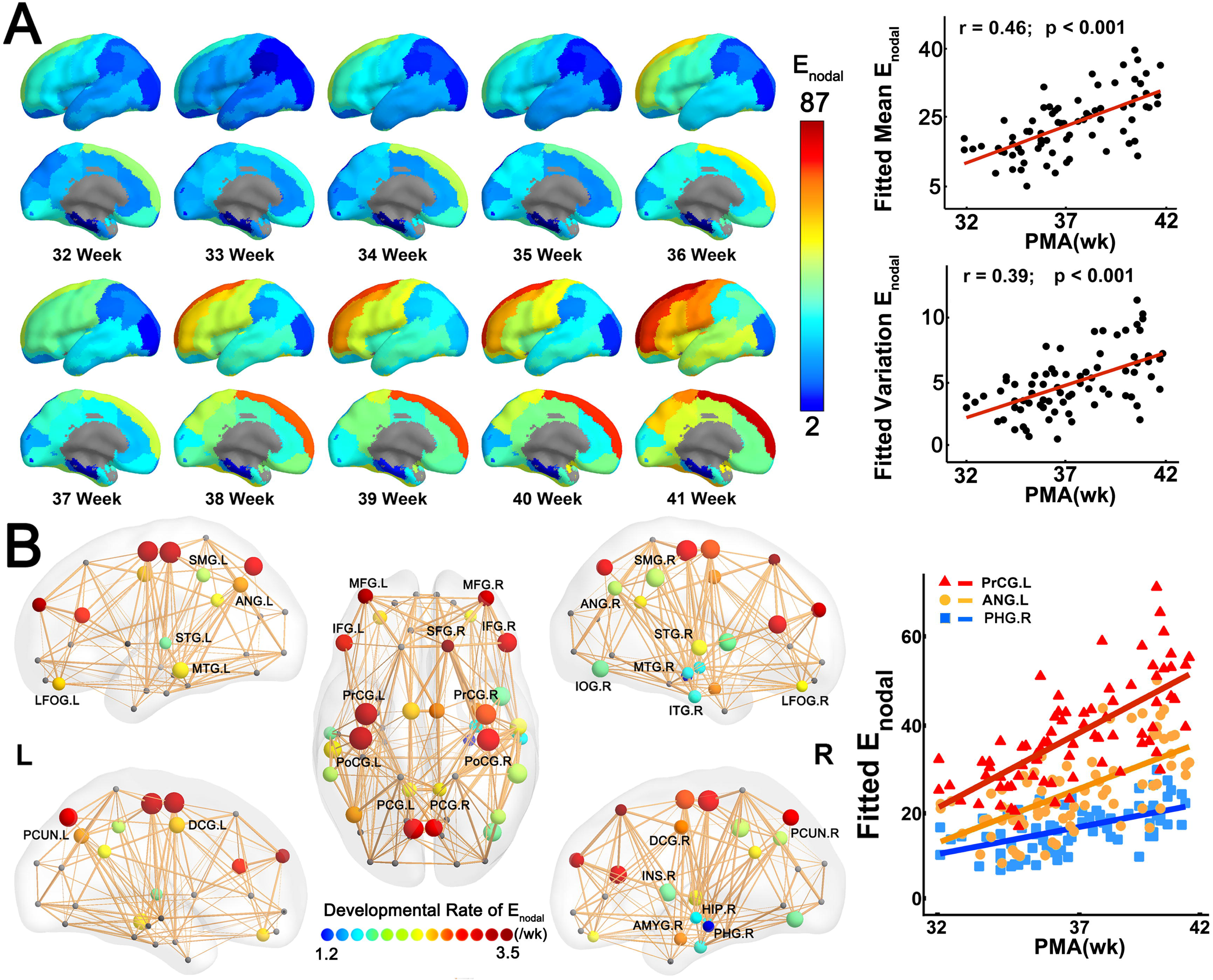
The heterogeneous development of nodal efficiency across brain regions. (A) On the left panel, fitted nodal efficiency maps at each week from 32 to 41 PMW demonstrate heterogeneous nodal efficiency distribution across the cortical surface. On the right panel, both mean and the standard deviation of nodal efficiency increased significantly with age. (B) On the left panel, 31 brain regions with significant and heterogeneous age-related increases of nodal efficiency are displayed as small spheres with colors (from blue to red) encoding different increase rates of nodal efficiency and sizes encoding the R values of the correlation between the nodal efficiency and age. The scatter plots on the right panel show significant age-related increases in nodal efficiency of three representative regions, namely, PrCG.L, ANG.L and PHG.R, from highest to lowest efficiency increase rate. The colors of the dots and the fitted lines for each representative region in the scatter plots are consistent with those encoding nodal efficiency increase rates shown on the left panel. Abbreviations: AMYG: amygdala; ANG: angular gyrus; DCG: dorsal cingulate gyrus; HIP: hippocampus; INS: insular cortex; IOG: inferior occipital gyrus; L/R: left/right; LFOG: lateral fronto-orbital gyrus; PCG: posterior cingulate gyrus; PCUN: precuneus; PHG: parahippocampal gyrus; PrCG/PoCG: precentral/postcentral gyrus; SFG/MFG/IFG: superior/middle/inferior frontal gyrus; SMG: supramarginal gyrus; STG/MTG/ITG: superior/middle/ inferior temporal gyrus.

#### Hub distribution, rich-club organization and faster nodal efficiency increases at the brain hubs

Figure 4A shows that the hub regions (red balls) are mainly distributed in the bilateral superior and middle frontal cortex, precentral and postcentral gyrus, superior parietal cortex and cingulate cortex. Consistent hub distribution across different PMW age groups from 31 to 41PMW was observed, as shown in Figure S2. Moreover, a characteristic rich-club organization with the normalized RC exceeding 1 (*ϕ*_norm_ = 1.21) was found for the backbone network, revealing a densely connected component between hub regions of the neonate structural connectome. Based on a data-driven two-cluster model for categorizing nodal efficiency, among 31 brain regions (shown in Fig. 3B) with significant age-dependent nodal efficiency increases, all 11 cluster-1 brain regions (blue circles in Fig. 4A and listed in Table 2) were part of the 16 hub regions (red balls in Fig. 4A) of the neonate connectome, indicating distinctively higher rate of efficiency increases during 31-42PMW at hub regions. Note that not all hub regions were characterized by statistically significant efficiency increases; however, those with significant efficiency increases were all cluster-1 brain regions. Fig. 4B shows significantly steeper efficiency increase trend line at the hub regions compared to non-hub regions (t = 6.85, interaction p < 10^-3^). Furthermore, by correlating the nodal efficiency increase rates and nodal efficiency measurements at 31 brain regions with significant age-dependent changes, we found significantly positive correlation between efficiency increase rate and efficiency measurements themselves (r = 0.78, p < 10^-3^) (Fig. 4C).

**Table 2.**
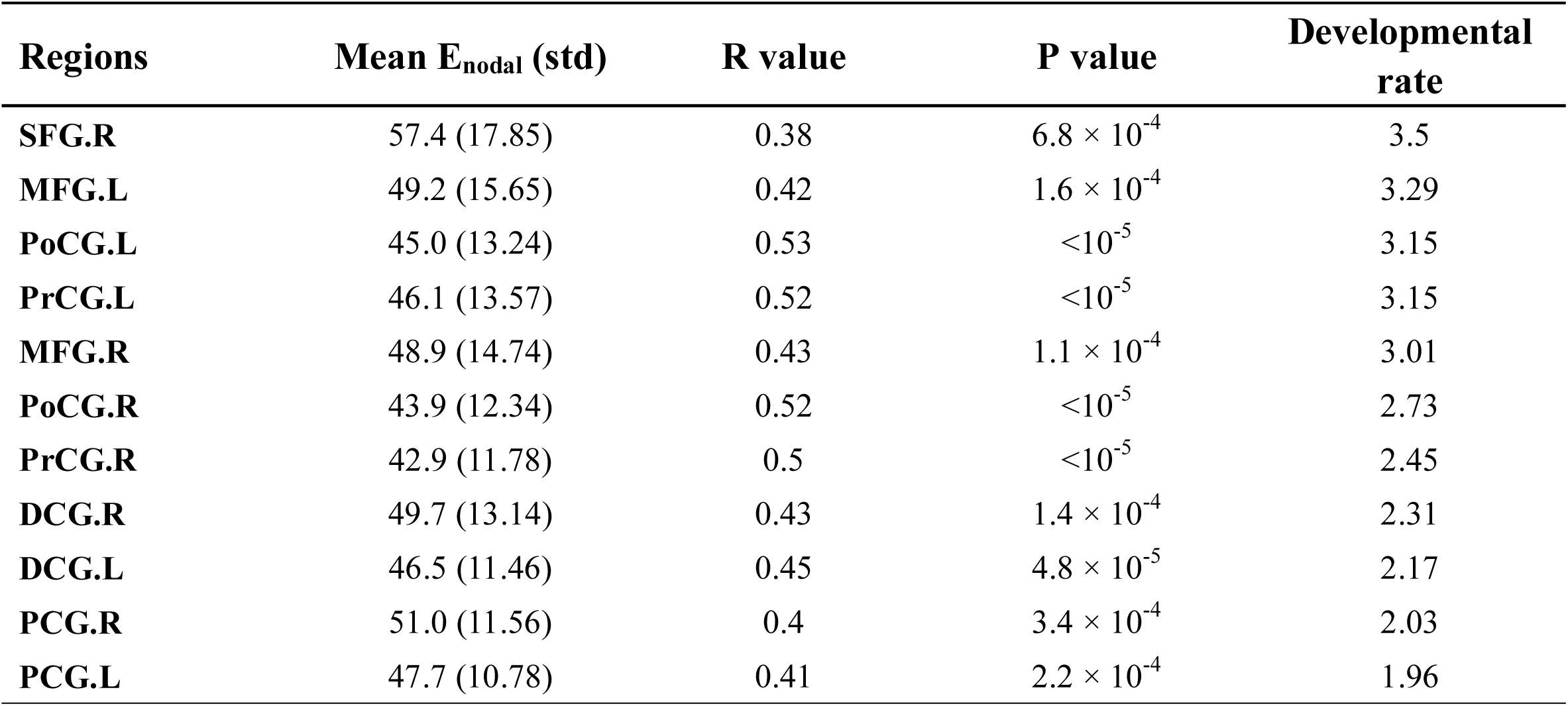
The brain hub regions with the statistically significant age-related increases in nodal efficiency. The hub regions were sorted by descending developmental rate. See legend of Figure 1 for abbreviation of brain regions.

**Figure 4.**
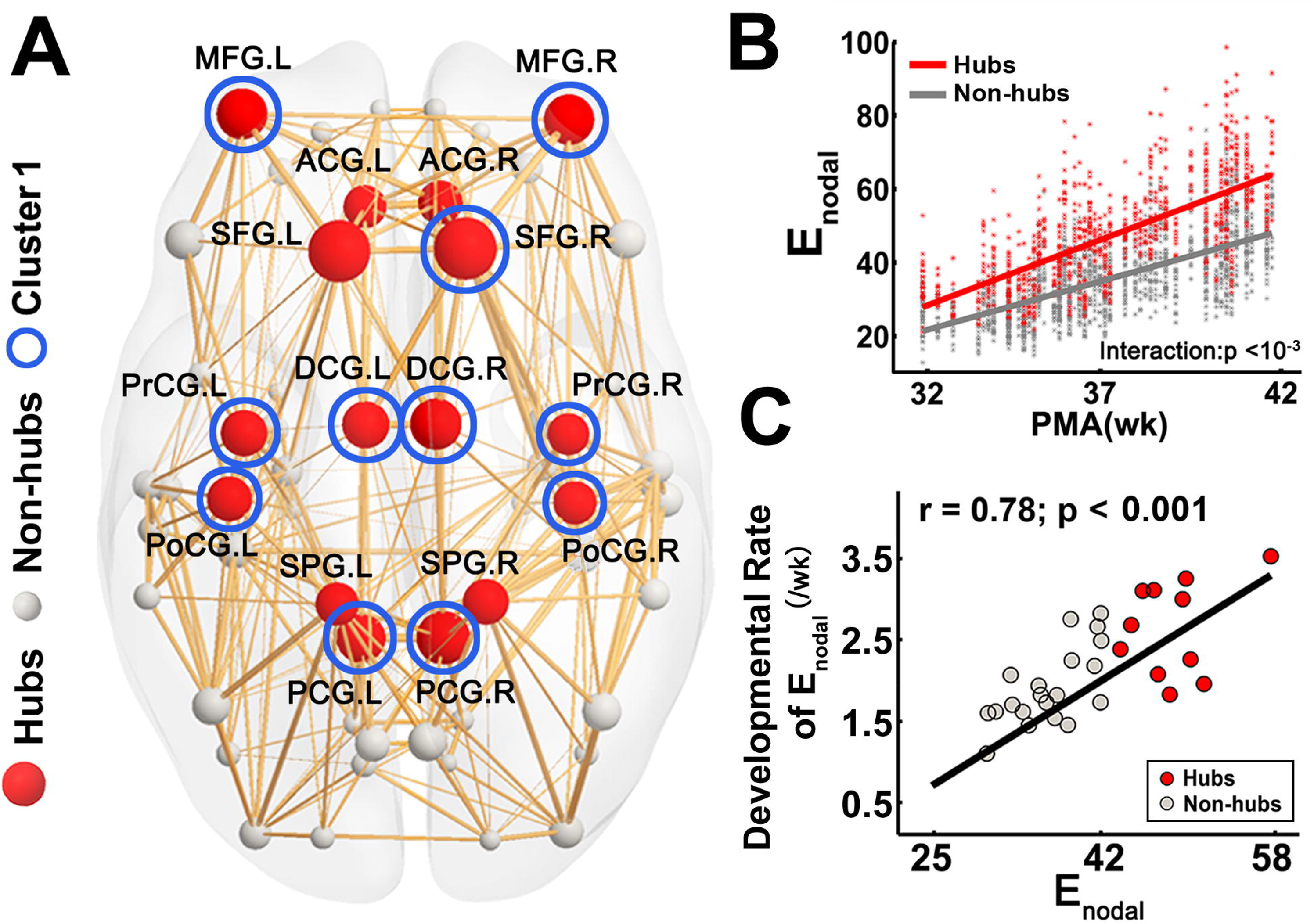
Hub distribution of the neonate connectome and higher developmental rates in hub regions compared to non-hubs. (A) A 3D representation of the hub distribution of the neonate structural connectome with the hub nodes in red and non-hub nodes in gray overlaid on the group-averaged backbone network. The size of the spheres encodes the averaged nodal efficiency across all neonates. Cluster-1 nodes identified by a data-driven clustering analysis were marked with blue circle. Of note, all cluster-1 nodes overlapped with the hub regions of the connectome. (B) Scatter plot showing more rapid age-related increases of nodal efficiency in hub regions than non-hub regions. The partial correlation between age and regional efficiency were fitted separately in all hub regions (red dots) and non-hub regions (gray dots). The interaction effect between age and hub category was significant (p <10^-3^). (C) Scatter plot showing significant linear correlation between the average nodal efficiency and developmental rate of nodal efficiency across the brain regions with significant nodal efficiency increases, with the hub regions in red and non-hub regions in gray. See legend of Figure 3 for abbreviations of brain regions.

### Faster edge strength increases in rich-club organization, in short-range connections and in intra-module connections

#### Faster edge strength increases in rich-club organization

NBS analysis revealed 101 significantly increasing edges (10% of all edges) connecting 56 nodes, which were widely distributed in bilateral frontal, parietal, temporal and limbic areas (Fig. 5A). For the connections with significant age-dependent changes, the rate of changes varied across the edges, reflected by differentially encoded edge width (Fig. 5A). Higher increase rates of edge strength were found in a few symmetric and short-range connections, including bilateral connections between superior and middle frontal gyrus, bilateral connections between precentral and postcentral gyrus, and bilateral connections between precuneus and posterior cingulate gyrus (Fig. 5A). With the connections classified into rich-club, feeder, and local connections (Figs 5A and 5B), significant differences in edge strength change rates were found among these three types of connections (Fig. 5B). Specifically, highest rate of edge strength increase was found in rich-club connections, followed by feeder and local connections (Fig. 5B).

**Figure 5.**
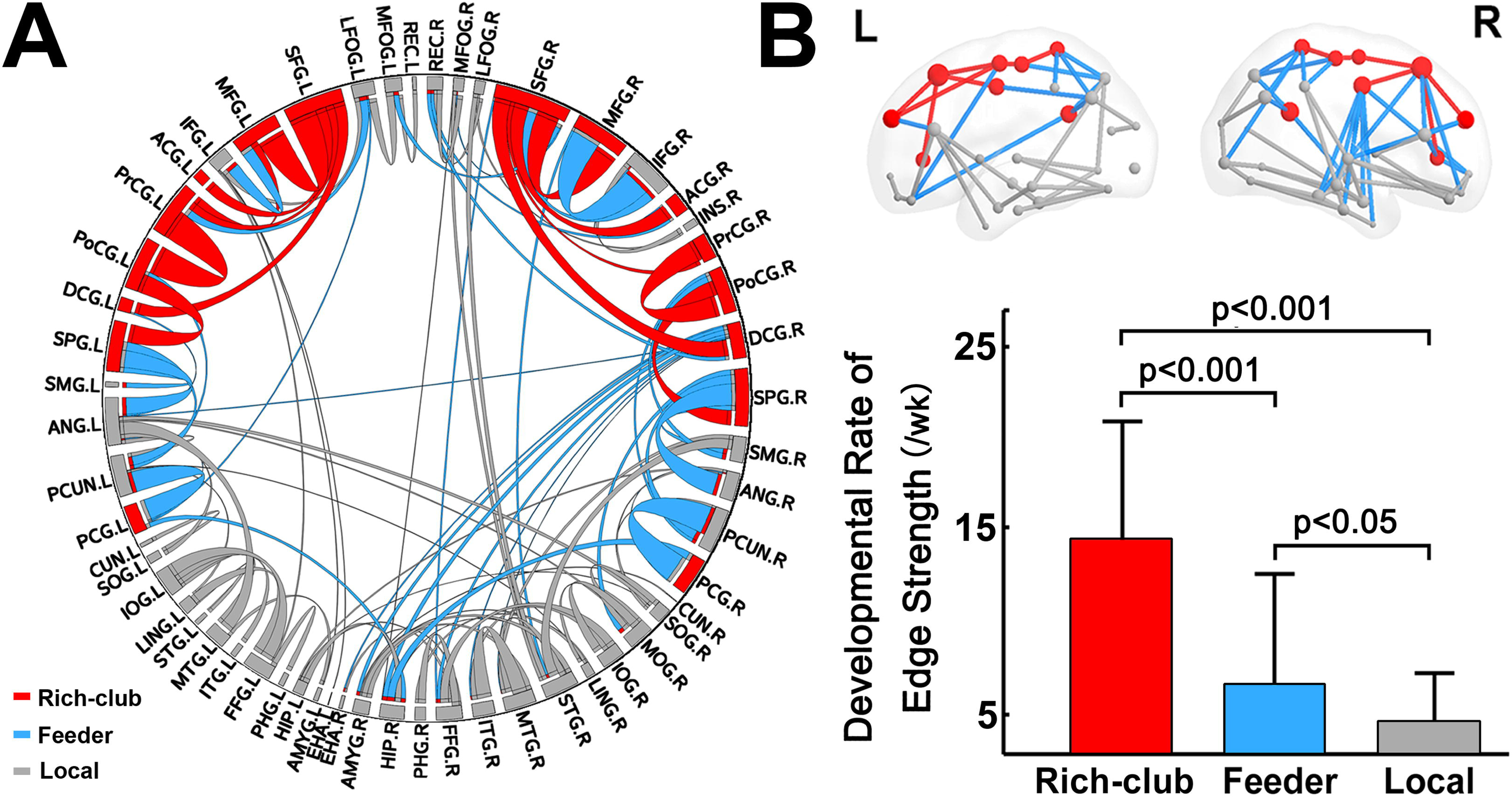
The components with significant age-related alterations revealed by NBS analysis and differential development rates of edge strengths in rich-club organization. (A) The NBS component is shown in a circle view with the color of the edges encoded by the categories of rich-club (red) feeder (blue) and local (gray) edges and size of edges encoded by the developmental rate. (B) The bar plot showing significant differences in edge strength developmental rate among the rich-club, feeder and local connections. See legend of Figure 3 for abbreviations of brain regions.

#### Faster edge strength increases in short-range connections

Representative short- and long-range connections based on the pathway length were demonstrated in Fig. 6A. Amongst short-range connections, rich-club (r = 0.37, p = 1.0 × 10^--3^), feeder (r = 0.49, p = 9.6 × 10^-6^) and local (r = 0.48, p = 1.0 × 10^-5^) connections (Fig. 6B) all increased significantly with age. By contrast, amongst long-range connections, only feeder (r = 0.24, p = 3.9 × 10^-2^) and local (r = 0.30, p = 8.7 × 10^-3^) connections increased significantly with age while no significant age-dependent changes were found for rich-club connections (r = -0.03, p = 0.75) (Fig. 6C).

**Figure 6.**
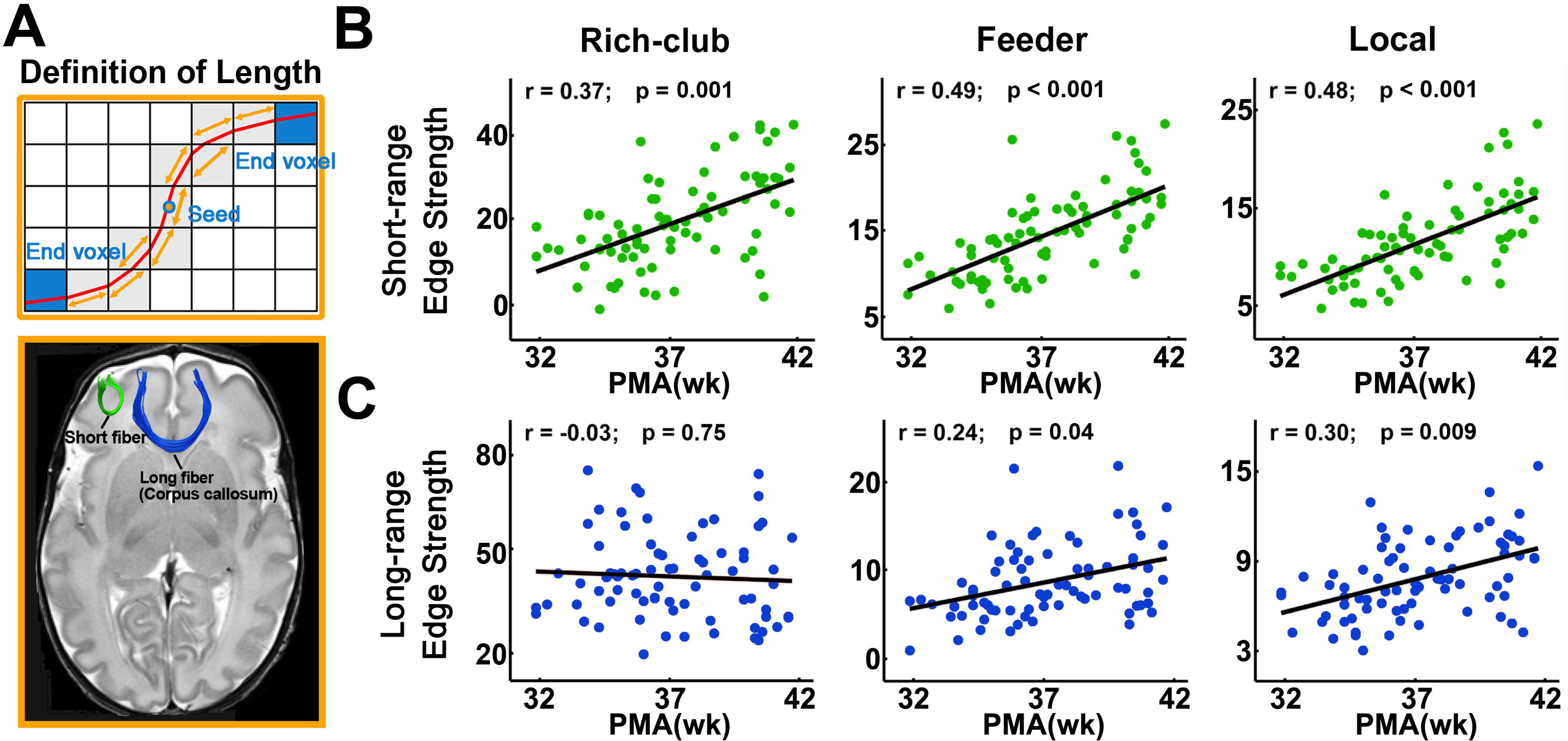
The development pattern of short- and long-range connections with age. (A) The definition of physical fiber length is shown in the upper panel. The physical length of a streamline reconstructed from deterministic tractography by following the main diffusion direction within each voxel was the length of the red curve. An illustration of short- and long-range fibers is presented in the lower panel. (B) Scatter plots showing significantly and relatively sharp age-related edge strength increases in different categories of short-range connections. (C) Scatter plots showing non-significant age-related changes of edge strength of rich-club long-range connections as well as significant but relatively mild age-related edge strength increases of feeder and local long-range connections.

#### Neonate brain modular organization and faster edge strength increases in intra-module connections

Significant modular organization was found in the structural networks of all individual subjects (Q > 0.34 for all subjects). No significant age-dependent alterations in the modularity and module number were found (all p > 0.05), suggesting that the modular organization remained stable during 32-41 PMW. Therefore, we only examined the modular parcellation of the group-based backbone network. A significant modular architecture of the backbone network was identified (Q_max_ = 0.42), separating the brain into five different modules (Fig. 7A, left panel). Hub regions were evenly distributed in different modules with provincial hubs located in the center of modules and connector hubs located in the boundaries. The bilateral precentral and postcentral gyrus and left superior and middle frontal gyrus were detected as provincial hubs, and the other hub regions were connector hubs (Fig. 7A). By comparing nodal efficiency increasing rates of these two types of hubs, we found the nodal efficiency increasing rate of provincial hubs are significantly higher than that of connector hubs (t = 2.91, p = 0.01) (Fig. 7B). In addition, we found that the age-dependent edge strength increase rate of within-module connections was higher than that of the between-module connections (t = 5.25, p = 0.003) (Fig. 7C). R values, p values and age-dependent edge strength increase rates from the correlation of edge strength and age for within-module and between-module connections are listed in the Supplemental Table S1.

**Figure 7.**
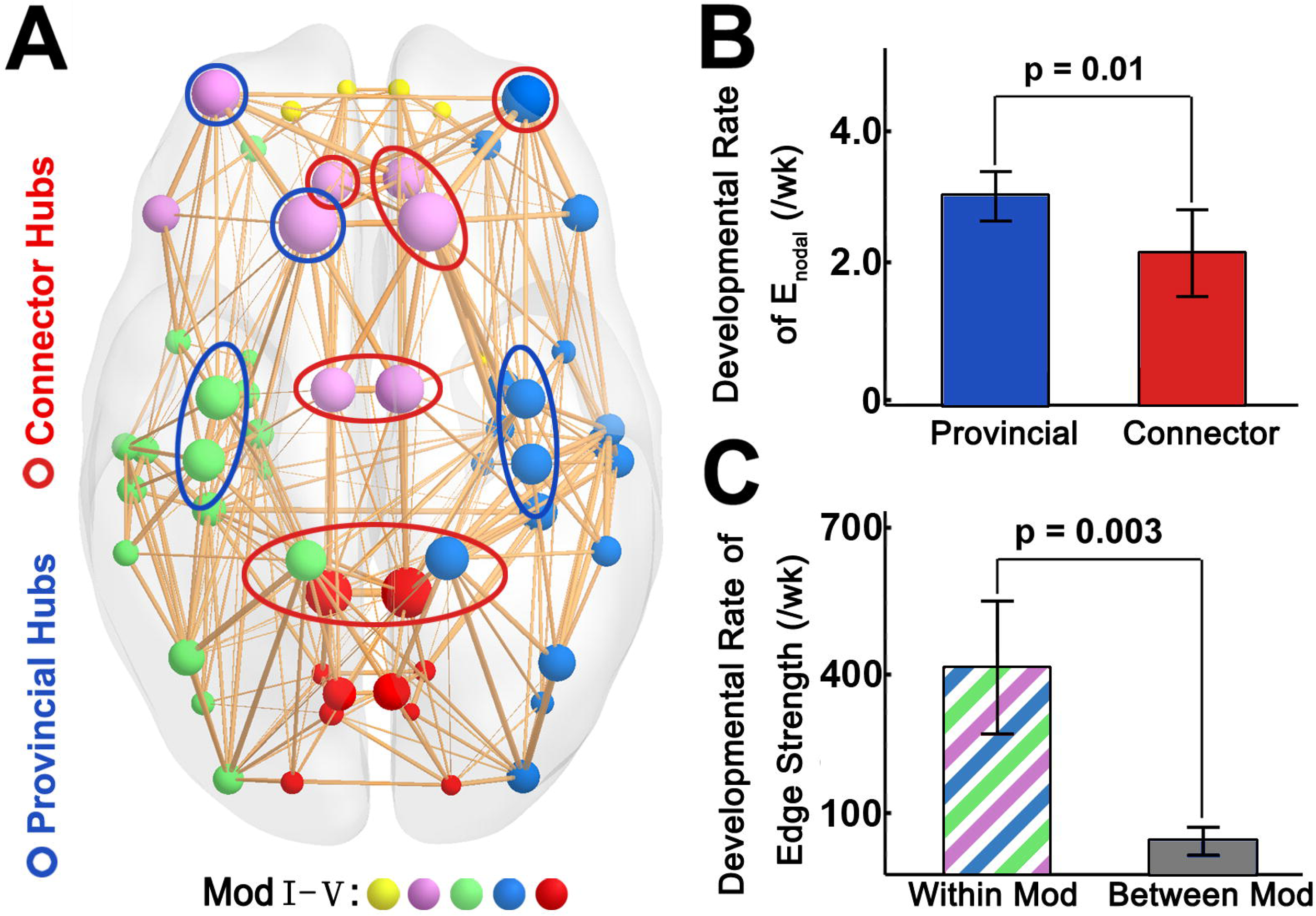
Higher developmental rates of edge strength of provincial hubs and within-module connections compared with connector hubs and between-module connections, respectively. (A) A 3D representation of the modular structure of the group-averaged backbone network with nodes in different colors corresponding to different modules and size encoding average regional efficiency. Module I was composed of bilateral orbital-frontal regions (yellow). Module II was composed of bilateral prefrontal regions (purple). Module □ mainly consisted of left pre/postcentral gyrus, temporal and superior parietal regions (green). Module □ mainly consisted of right pre/postcentral gyrus, temporal and parietal regions (blue). Module □ mainly consisted of bilateral posterior parietal regions (red). The provincial hubs and connector hubs were marked with blue and red circles, respectively. (B) The bar plot showing higher developmental rate of provincial hubs than that of connector hubs. (C) The bar plot showing higher developmental rate of within-module (Within Mod) connections compared with that of between-module (Between Mod) connections.

### Age prediction and reproducible findings with a high-resolution parcellation scheme

#### Age prediction

Fig. 8A shows that the postmenstrual ages in weeks of the neonates can be predicted by the nodal efficiency of brain structural connectome, with a correlation r = 0.76 between the actual and the predicted postmenstrual age. It is noteworthy that hub regions showed higher prediction accuracy (r = 0.81) (Fig. 8B) than non-hub regions (r = 0.66) (Fig. 8C), suggesting a stronger association between hubs and postmenstrual age as compared to non-hubs.

**Figure 8.**
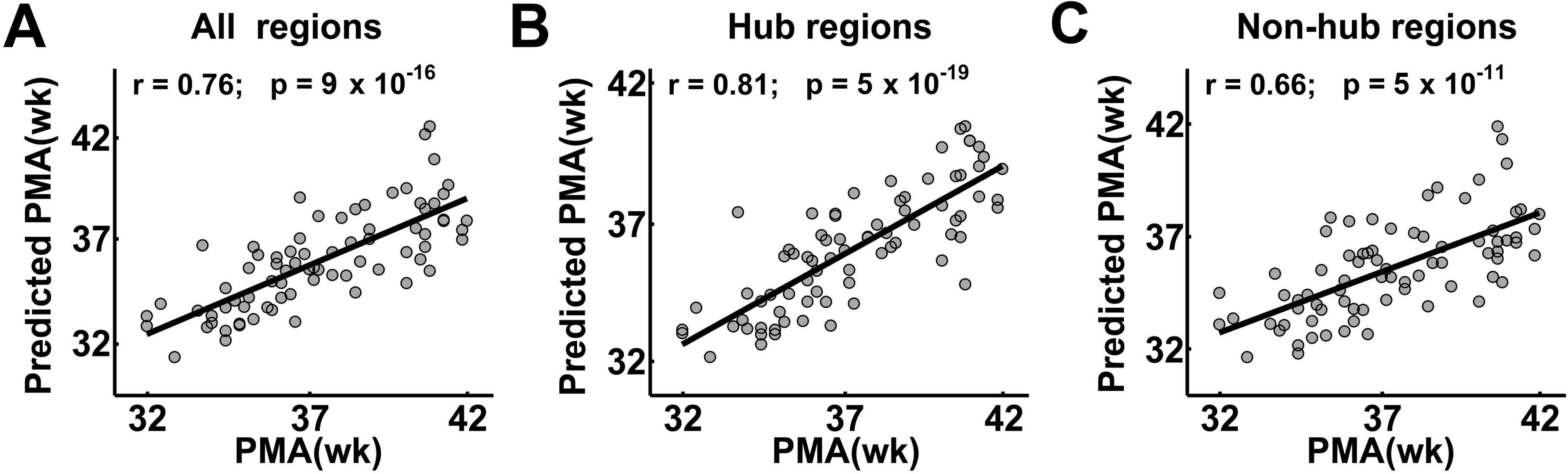
The prediction of individual age based on nodal efficiency of all brain regions (A), hub regions (B) and non-hub regions (C). The scatter plots depict actual versus predicted age. Pearson correlation coefficient between the actual and predicted ages are shown to assess the prediction accuracy.

#### Reproducible findings with a high-resolution parcellation scheme

Similar age-related development trend lines of global and regional network properties were observed when the analyses were repeated utilizing the higher resolution parcellation. This included significant age-dependent increases in global and local network efficiency (Eglob: r = 0.58, p = 5.9 × 10^-8^; Eloc: r = 0.63, p = 1.2 × 10^-9^) (Fig. 9B). Likewise, the brain regions with most rapid age-dependent increases of nodal efficiency were distributed in the precentral and postcentral gyrus cortex and posterior parietal cortex (Fig. 9C), consistent with the findings from low-resolution parcellation. Similar hub regions were found mainly located in the bilateral orbito-frontal cortex, bilateral precentral and postcentral cortex, bilateral superior parietal cortex and temporal cortex (Fig. 9D). Finally, similar to Fig 4C, a significant positive correlation between mean nodal efficiency and their developmental rate was also observed (r = 0.43, p = 9.0 × 10^-12^) in regions with significant age-related alterations in high-resolution networks (Fig. 9E). These results jointly indicated that the maturation patterns of the neonate connectome were largely independent of the cortical parcellation schemes.

**Figure 9.**
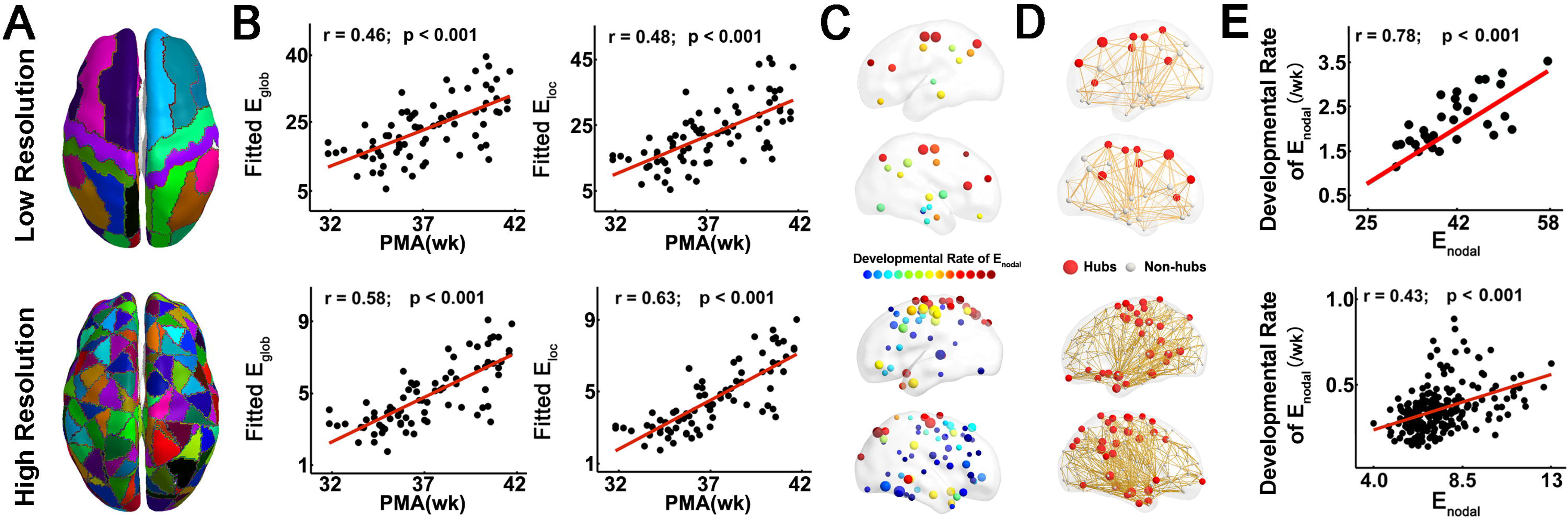
Reproducible age-dependent alterations with a high-resolution parcellation scheme. (A) The parcellation with JHU-58-region neonate atlas (Oishi et al. 2011) and the high resolution cortical parcellation with 256 ROIs. (B) Scatter plot showing significantly linear increases with age for the global efficiency and local efficiency in both low- and high-resolution networks. (C) Region distributions with significant age-dependent changes of nodal efficiency for low- and high-resolution network displaying as small spheres with colors encoding the developmental rates of nodal efficiency and sizes encoding the R values. (D) Similar hub distributions between low- and high-resolution networks with the hub nodes in red and non-hub nodes in gray and size encoding average regional efficiency across all neonates, overlaid on the group-averaged network backbone. (E) Scatter plots showing significant linear correlation between the average nodal efficiency and developmental rate of nodal efficiency across the brain regions, in both low- and high-resolution networks.

## Discussion

Using connectomic analyses of dMRI images with relatively high resolution (1.5 × 1.5 × 1.6 mm^3^) from 77 preterm and term-born neonates scanned at 31 to 42 PMW, we found rapid and regionally differentiated maturation with faster connectivity increases taking place mainly at the brain hubs and rich-club, especially for the short-range and within-module connections, resulting in a more segregated structural connectome near term-equivalency. The brain hubs with faster age-dependent nodal efficiency increases are distributed in primary sensorimotor regions, superior-middle frontal and posterior cingulate gyrus, while the hub distribution remains almost unchanged during 31-42PMW. The faster connectional maturation at these hub regions was supported by a data-driven cluster analysis. Compared to long-range or between-module connections, short-range and within-module connections appeared to develop more rapidly during 31-42PMW, contributing to emergence of the rich-club organization and brain modules. Efficiency measures of all brain regions, especially those of hub regions, accurately predicted neonatal age in PMW. The findings in this study shed light on the spatiotemporal principles of brain connectome development during this critical period, offering references for aberrant brain organization that may be associated with neurodevelopmental disorders. The highly accurate prediction of age at the identified hubs suggests that these core regions may serve as biomarkers indicating the ontogeny of early brain development. Collectively, the results revealed rapid increases of the hub and rich-club connections, resulting in structural segregation that underlies functional segregation (Cao et al. 2017a) and emergence of certain primary brain functions during the same developmental period.

### Segregation of neonate brain structural connectome

As shown in Figure 2, dramatic global and local efficiency increases during 31-42PMW were found with global topology analysis, suggesting that the white matter maturation contributes to a more topographically efficient and compact network during this critical period, consistent to the existing literature (van den Heuvel et al. 2015). The increases of regional efficiency of structural network are widespread in Figure 3, contributing to global efficiency increases in global topology analysis. Despite overall increases, the pattern of age-dependent nodal efficiency increases is not uniform across the brain regions (Fig. 3), with some regions demonstrating rapid increases in nodal efficiency while other regions remain almost unchanged, contributing to the segregation of baby brain connectome during 31-42 PMW. It is noteworthy that the regions with higher nodal efficiency exhibited higher developmental rate too (Fig. 3). Figure 4 further demonstrated that among all the nodes with significant age-dependent efficiency increases, fastest efficiency increases coincided with the brain hubs. More accurate prediction of individual age was found at the hub regions of the structural connectome than other regions (Fig. 8). These results suggest that selective strengthening of hubs is prominent during the last several weeks before normal time of birth. On the other hand, the hub distribution across the ages during 31-42PMW are almost unchanged (Supp Fig. 2). This supports that the increased segregation of the developing connectome is achieved by increasing the connectivity to and from key hubs, established early in gestation, rather than by altering of hub distribution. Brain hubs occupy a dominant position in information transfer (Xu et al. 2010) and have higher levels of metabolic energy consumption and higher rates of cerebral blood flow than peripheral nodes (Liang et al. 2013; Tomasi et al. 2014). The rich club of hub regions observed here in bilateral precentral and postcentral gyrus, posterior cingulate gyrus, superior and middle frontal cortex, are consistent with prior observation in neonates (Ball et al. 2014; Pandit et al. 2014; van den Heuvel et al. 2015). Moreover, these hubs show a strong correspondence with those of the adult structural connectome (Hagmann et al. 2008; Gong et al. 2009; van den Heuvel and Sporns 2011). Our findings and previous studies jointly suggest that these network hubs are not only critical for the neonatal brain to get ready for postnatal neural growth, but also play a key role in organizing the connectome throughout brain development. In addition, functional network segregation was observed in a subset of the same preterm cohort by analyzing resting-state fMRI dataset (Cao et al. 2017a). The structural network segregation in the present study is likely to underlie the functional segregation.

Higher increase rates of edge strength in rich-club edges were found in short-range connections, compared to those in long-range connections (Fig. 6). Moreover, higher developmental rates were found in provincial hubs than connector ones (Fig. 7B) and in within-module connections than between-module ones (Fig. 7C), with the module distribution of the neonate brains at 31-42PMW (Fig. 7A) similar to that of adult brains (Hagmann et al. 2008). The edges within modules and provincial hubs mainly contribute to connections within particular systems, as compared to global integration. Faster increases of edge strength and nodal efficiency in particular systems make the network more specialized and segregated during maturation. These results are consistent to general understanding of normal developmental course of structural network characterized by gradual maturation from local and proximity-based connections supporting primary functions to a more distributed and integrative topology supporting higher cogntive functions (Hagmann et al. 2010; Yap et al. 2011; Bullmore and Sporns 2012; van den Heuvel et al. 2012; Tymofiyeva et al. 2013; Collin et al. 2014; Vertes and Bullmore 2015).

### Differentiated maturation of brain regions with higher rate of efficiency increases at the brain hubs

The nodal efficiency is heterogeneously distributed and the increase rate of the nodal efficiency is also differentiated across the brain regions from Figure 3. Among all brain nodes with statistically significant increases of nodal efficiency, higher rates of nodal efficiency increases were found at certain brain hubs (Fig. 4). From Table 2, these hub regions with significant nodal efficiency changes are the left and right precentral and postcentral gyrus, left and right dorsal cingulate gyrus and posterior cingulate gyrus, as well as some frontal gyri. As elaborated below, these regions were consistently found to play an important role in early brain development. The left and right precentral and postcentral gyrus are essential for primary sensorimotor functions. Previous functional connectivity studies (Doria et al. 2010; Smyser et al. 2010; Fransson et al. 2011) found these regions among the earliest appearing functional networks identified from resting-state fMRI. Moreover, the identified hubs of neonate functional connectome are largely confined to primary sensorimotor regions (Fransson et al. 2011; Gao et al. 2011; Cao et al. 2017a), which distinguishes the neonate brain from adult brain. Considering that this differential pattern is observed across modalities, including PET (Chugani et al. 1987; Chugani 1998), it is likely that early maturation of receptive sensory areas may not only be helpful for the basic survival functions at birth (Buckner and Krienen 2013) but also support the later maturation of higher order and multimodal integrative areas (Guillery 2005). Left and right posterior cingulate gyri, as a functional core of the default-mode-network (Fransson and Marrelec, 2008), are also regions with high rates of nodal efficiency increases (Table 2). The higher rate of efficiency increases of these gyri may underlie in infancy the emergence of primitive default-mode network (Gao, et al., 2009) which is critical for the neonates to develop a sense of self (e.g. Uddin et al. 2007). Observation of the most significant nodal efficiency increases in the hub regions of primary sensorimotor region and posterior cingulate gyrus offers the structural connectivity basis for the coupling of structural and functional topology development in the baby brain connectome at these regions. Other hubs with significant nodal efficiency increases include middle and superior frontal gyri (Fig. 4 and Table 2). The higher rates of connection increases in frontal lobe during preterm development have been observed in recent structural connectivity studies (Brown, et al., 2014; Pandit et al., 2014). Active frontal cortical maturation has also been reflected by cortical FA changes. Sharp decrease of frontal cortical FA, a measure quantifying the dendritic arborization in the cerebral cortex, has been consistently found in several studies using cortical FA to delineate the cortical microstructural developmental pattern of the preterm brains (DeIpolyi, et al., 2005; Ball et al., 2013; Yu et al., 2016). The observation of structural connectivity hubs at superior and middle frontal cortex could be related to active cortical microstructural activities in these regions.

### Emergence of rich-club organization in early developing brain

Figure 5 shows rich-club organization (van den Heuvel and Sporns 2011) consisting of all hubs exhibited in Figure 4A. In the present study, rich-club organization was found in the structural connectome of preterm brain at the age of as early as 31PMW. Despite that the early emergence of rich-club organization has also been revealed in recent studies of neonate structural connectome (Ball et al. 2014; van den Heuvel et al. 2015), the present study revealed the developmental rate of rich-club and other network properties by employing the neonate dataset with relatively large sample and evenly distributed ages from 31-42 PMWs, offering a new insight into the asynchronous development across brain regions. Heterogeneous increases of nodal efficiency among brain regions and faster increases at the brain hubs may drive the emergence of rich-club organization in the period of 31-42 PMW. Rich-club organization was also contributed by more rapidly strengthened edges in rich-club connections than those in other connections, as demonstrated by fastest edge strength increases in rich-club connections, followed by the feeder connections and the local connections (Fig. 5B). The “rich-get-richer” principle of network evolution (Barabasi and Albert 1999) means that new connections are preferentially associated to the nodes with many connections and is reflected by the findings of more rapid growth of hubs and rich-club connections in the present study. Additionally, the accelerated growth of hub regions leads to a wider degree distribution including more highly-connected hubs as well as more non-hub nodes with few local connections (Kaiser 2017), possibly underlying important functional roles of hub regions after birth and functional segregation of brain regions.

### Short-range and long-range connections

As can be observed from Figure 5, most of significantly increasing edges were short-range connections. The present study revealed that faster increasing edge strength of short-range connectivity compared to that of long-range connectivity may facilitate the structural connectome segregation process centered at rich-club organization (Figs. 5 and 6) during 31-42PMW. Developmental rates of short-range cortical-cortical connections in rich-club edges were higher than those of long-range connections (Fig. 6). Particular growth of short rich-club edges may also enhance local neuronal operations and segregation of modules, supported by modular analysis shown in Figure 7. Delineation of the brain connectivity pathways with dMRI has revealed white matter morphological dynamics from early 2^nd^ trimester to birth (Huang et al. 2006; Huang et al. 2009; Takahashi et al. 2012; Ouyang et al. 2015). Among all white matter tract groups, the long-range association tracts connecting cortical regions are those emerging relatively late. For example, the arcuate fasciculus, key for the language function development, is not well developed until 2 years of age (Zhang et al, 2007). Other connectomic studies have found that the increasing edges in prenatal and preterm developmental stage consisted of many short local connections and limited long-range connections (Takahashi et al. 2012; Brown et al. 2014). Faster increasing short-range connections during 31-42PMW may constitute the pivotal edges of the rich-club backbone and mediate specialized functional process in local integration (Park and Friston 2013).

### Limitations, technical considerations and future directions

We tested the effects of different cortical pacellation schemes and found that the maturation patterns of the neonate connectome were largely independent of the cortical parcellation schemes, as demonstrated by Figure 9. Besides effects of different parcellation schemes on connectomic analysis results, several issues need to be further considered for future studies. First, the dataset used in this study was obtained using a cross-sectional design. Future studies with longitudinal design may need be considered to eliminate the effects of individual differences, despite that the age-related structural connectomic changes are dominant in this very dynamic early developmental period. Second, deterministic tractography was used for the reconstruction of WM tracts, which may have resulted in the loss of existing fibers due to the “fiber-crossing” problem (Mori and van Zijl 2002). The tractography techniques more robust to fiber-crossing, such as probabilistic tractography (Behrens et al. 2007), can be considered to define the network edges in future studies. Third, it has been found that preterm birth was associated with altered microstructural development (e.g. Boardman et al. 2010; Rathbone et al. 2011) and adverse neurodevelopmental outcomes (Woodward et al. 2006). Despite preterm birth effects, MRI examinations of preterm infants have been predominantly used to understand brain development during the 3^rd^ trimester. Several other studies using dMRI also indicated dramatic reconfiguration during the last 10 weeks prior to normal time of birth (e.g. Ball et al. 2013; Brown et al. 2014; van den Heuvel et al. 2015). Exposure to the extrauterine environment could constitute part of the observed network reorganization (Karolis et al. 2016; Batalle et al. 2017), but these effects would be relatively subtle compared with effects of very dynamic development during 3^rd^ trimester (Bourgeois et al. 1989; Kostović 1990). Nevertheless, it is likely that the disruption of the network could become apparent in years subsequent to premature birth. Recent advances of in-utero MRI (e.g. Kasprian et al. 2008; Thomason et al. 2013; Mitter et al. 2015; van den Heuvel and Thomason 2016) could alleviate the preterm effects. Fourth, the segregation has also been found in functional connectome development during this period in our previous study (Cao et al. 2017a). It is noteworthy that the cohort of functional connectome study (Cao et al. 2017a) was a subset of the cohort used in the present one. With the same segregation processes found in the structural connectome, the mechanistic relationship on how structural connections underlie functional ones has yet to be delineated. Recently developed approach, such as the one used for understanding network level structure-function relationships (Mišić et al. 2016), could offer the insights of mechanistic structure-function relationship in this specific developmental period. Finally, further studies on relationship of maturation of hubs or rich-clubs at certain brain regions and emergence of the brain functions could contribute to understanding of general developmental principle.

## Acknowledgments

This study was sponsored by NIH (Grant Nos. MH092535 and MH092535-S1, HH), the 973 program (Grant No. 2013CB837300, NS), the National Natural Science Foundation of China (Grant Nos. 81471732, 81671761, NS; 81628009, HH), the Fundamental Research Funds for the Central Universities (Grant No. 2017XTCX04, NS), and the Interdisciplinary Research Funds of Beijing Normal University (NS).

## Supplemental Figures

**Figure S1.** Age-related changes of the preterm brain global network metrics, Lp (shortest path length), Cp, Gamma (normalized Cp) and Lamda (normalized Lp). Significant decrease of Lp with age and non-significant changes of Cp, Gamma and Lamda with age are demonstrated in the scatter plots.

**Figure S2**. Almost unchanged brain hub distributions across different age groups, 31-34, 34-36, 36-38, 38-40 and 40-42 PMW (week). The distribution is represented in 3D with the hub nodes in red and non-hub nodes in gray overlaid on the group-averaged backbone network. The size of the spheres encodes the nodal efficiency normalized in a given age range.

